# Evaluating ultrastructural preservation quality in banked brain tissue

**DOI:** 10.1101/2025.05.09.652503

**Authors:** Macy Garrood, Alicia Keberle, Allison Sowa, William Janssen, Emma L. Thorn, Claudia De Sanctis, Kurt Farrell, John F. Crary, Andrew T. McKenzie

## Abstract

The ultrastructural analysis of postmortem brain tissue can provide important insights into cellular architecture and disease-related changes. For example, connectomics studies offer a powerful emerging approach for understanding neural circuit organization. However, electron microscopy (EM) data is difficult to interpret when the preservation quality is imperfect, which is common in brain banking and may render it unsuitable for certain research applications. One common issue is that EM images of postmortem brain tissue can have an expansion of regions that appear to be made up of extracellular space and/or degraded cellular material, which we call ambiguous interstitial zones. In this study, we report a method to assess whether EM images have ambiguous interstitial zone artifacts in a cohort of 10 postmortem brains with samples from each of the cortex and thalamus. Next, in matched samples from the contralateral hemisphere of the same brains, we evaluate the structural preservation quality of light microscopy images, including immunostaining for cytoskeletal proteins. Through this analysis, we show that on light microscopy, cell membrane morphology can be largely maintained, and neurite trajectory visualized over micrometer distances, even in specimens for which there are ambiguous interstitial zone artifacts on EM. Taken together, our analysis may assist in maximizing the usefulness of donated brain tissue by informing tissue selection and preparation protocols for various research goals.

## Introduction

Since the 1950s when the synapse was first visualized with an electron microscope, the ultrastructural analysis of brain tissue has enabled many critical discoveries about the structure of the brain (Sotelo, 2020; Nahirney and Tremblay, 2021). In recent years, EM-based connectomics in particular has enabled key insights into larger-scale neural circuitry across diverse species (Lichtman et al., 2014; Shiu et al., 2024). Expanding connectomics to a larger set of human brains may allow us to answer fundamental questions about our brains in health and disease. However, data collection for connectomics relies on high-quality tissue preservation that enables the reliable tracing of neuronal processes and identification of synaptic connections. These requirements are challenging to achieve in human brain tissue samples, which generally have quality limitations due to agonal factors, the postmortem interval (PMI), and impediments in preservation methods (McKee, 1999; McFadden et al., 2019; McKenzie et al., 2022). To date, most volume electron microscopy studies have been performed on non-human brain tissue (Chiappini et al., 2025). The successful volume electron microscopy and connectomics studies on human brain tissue have been limited to tissue that is selected for having ideal preservation quality, such as surgical biopsy tissue or deceased donor tissue with very low PMI (Shapson-Coe et al., 2024; Plaza-Alonso et al., 2025).

While mapping an entire human brain with electron microscopy far exceeds our current technological capabilities, the ability to image nanoscale neural connectivity in tissue from a wider set of banked brains would still be valuable for several reasons. First, brain banks serve diverse research communities, and investigators require access to tissue from different brain regions and with different donor characteristics, which may not always be possible to acquire from surgical biopsies. Second, larger sample sizes are critical for statistically robust inferences regarding the structural correlates of disease, as the history of genomics has shown (Uffelmann et al., 2021). Third, advances in imaging and computing technologies are rapidly improving our ability to analyze neural circuits at scale. As these tools continue to develop, having access to more banked brain tissue whose ultrastructure can be profiled may enable increasingly larger scale connectomics studies of the human brain. Given these considerations, there is a critical need to investigate whether current techniques used in brain banking are sufficient to maintain the structural integrity needed for future connectome analyses, despite the fact that our present connectome imaging and reconstruction capabilities are still limited.

There is a lack of standardized methods to assess brain preservation quality, which means that it may be useful to consider first principles (McFadden et al., 2019; Wahyudi et al., 2025).

Structural changes can occur prior to fixation, during fixation, and during subsequent tissue processing, and may include membrane blebbing, vacuoles, cytoplasmic washout, and chromatin alterations (Garman, 2011; Krassner et al., 2023; McKenzie et al., 2024). Some of these artifacts, such as blebbing or vacuoles, may not prevent us from tracing the connectome, instead only leading to the expansion or compression of certain cellular structures, which can still be adequately visualized. A more significant challenge arises from the observed phenomenon in some samples of enlarged, electron-lucent, non-membrane-bound regions between cells, which we call ambiguous interstitial zones (AIZs). Crucially, these AIZs are usually also associated with an apparent decrease in the density of visualized cellular structures, such as thin neurites. Instead, the AIZs often contain what appears to be poorly defined structures, possibly cellular debris. These poorly defined structures are not usually visible in tissue that has been preserved with a negligible duration of ischemia prior to preservation. These AIZs and associated poorly defined structures have been described in previous literature, for example as causing a “lacey” appearance in postmortem brain tissue (Palay et al., 1962; Liewald et al., 2014; Lewis et al., 2019).

Mechanistically, very early in the PMI, the extracellular space is actually expected to shrink due to ischemic cell swelling (Nicholson and Syková, 1998). As the PMI becomes prolonged, biomolecular structures break down and cell membrane integrity is eventually lost, leading to passive fluid redistribution and an expected expansion of the apparent extracellular space (Krassner et al., 2023). This excess fluid may affect fine neural structures in various ways, from complete dissolution in severe cases, to preservation of some elements but with insufficient compactness for visualization in others. The difficulty in visualization may be in part because microscopy depends on biomolecules being densely aggregated enough to maintain their structural integrity during tissue processing and bind adequately with staining chemicals (Wang and Minassian, 1987). However, during prolonged PMI, these biomolecular networks progressively dissociate, reducing their density.

Previous studies have found that fixation and sample preparation methods can influence the visualized ultrastructure of brain tissue (Schiff and Gennaro Jr., 1979; Small, 1981; Sele et al., 2019; Eberhardt et al., 2022; Shafiei et al., 2024). For example, one study reported that there can be significant extraction of proteins and lipids during EM preparation, creating large empty spaces, which is an issue that worsens with longer PMI (Lewis et al., 2019). They found that adding 0.1% glutaraldehyde to their formalin fixative partially rescued this. Another study found that the use of embedding techniques for EM at room temperature, which is standard for the field, introduced ultrastructural artifacts such as swollen mitochondria, disrupted membranes, and extraction of cellular components (Sosinsky et al., 2008). However, these artifacts were avoided when fixation was instead followed by a high-pressure freezing protocol, which led to images with smoother cell membranes and more dense cytoplasm. Finally, one study tested more than one hundred protocol variations on human brain tissue and found that even subtle differences in fixation, washing, dehydration, and embedding can affect the ultrastructural appearance, including the preservation of extracellular space (Karlupia et al., 2023).

Our working hypothesis is that the observed artifacts on ultrastructural imaging may result solely from structural degradation due to decomposition prior to fixation, and/or may be partially accounted for by our current methods of sample preparation, staining, and imaging, which are not optimized for decomposed tissue. If true structural loss is the dominant factor, then strict constraints on PMI and preservation procedures may be necessary for selecting samples that are suitable for connectome imaging. On the other hand, if enough morphomolecular markers are still theoretically present to reconstruct the cellular skeleton but are not visualized with current methods due to insufficient optimization for this purpose, then improved postfixation, embedding, staining, imaging, and analysis methods could potentially recover them.

In this study, we attempt to address the challenge of distinguishing true structural degradation from visualization artifacts in postmortem brain EM images. We develop a metric to assess the extent of AIZs in the EM images as an index for evaluating preservation quality. We also analyze matched samples from the contralateral hemisphere stained with hematoxylin and eosin (H&E) and via immunohistochemistry, to attempt to identify cases where cellular structures remain intact on light microscopy. Our goal is to help improve our methods for assessing ultrastructural preservation quality and explore potential discrepancies between preservation quality at the light and electron microscopy levels.

## Methods

### Brain banking procedures

Anatomical whole body donations were performed by a partner whole body donation organization operating under Oregon Health Authority regulations. Additionally, through our canine brain bank program, we were donated the body of one deceased canine following euthanasia by a licensed veterinarian, with signed owner consent for research use (Sándor et al., 2021). The Apex Neuroscience Brain and Tissue Bank operates under an exemption determination issued by the Pearl Institutional Review Board (IRB) after the submission of our protocols for review.

The PMI was calculated as the time elapsed between death and the initiation of the preservation procedure. When the day of death but not the hour was known, the time of death was estimated at the middle of the day (12 pm) to provide a standardized approach for PMI calculation in cases with incomplete time data.

One of the brain specimens (donor #57) was fixed by immersion in 10% neutral buffered formalin (NBF, Azer Scientific NBF55G) alone. The rest of the human brains were perfused *in situ* with the use of a peristaltic pump, following cannulation of the bilateral carotid arteries with 10% NBF. For several of the donors (donors #7, 30, 34, and 37), the carotid arteries were accessed via dissection in the anterior cervical region. For the remaining human donors, the cephalon was isolated via dissection at the approximate spinal level of C4-C5 (Turkoglu et al., 2014). This allowed for cannulation of the bilateral internal carotid arteries and clamping of the vertebral arteries to prevent outflow through them. The one canine brain was from a 2.3 kg Miniature Pinscher whose brain was perfused *in situ* with 10% NBF via cannulation of the left ventricle and the use of a peristaltic pump.

The brains were removed from the skull following standard procedures (Adams and Murray, 1982). They were then immersed in 10% NBF at 4°C for at least one month prior to further processing (McKee, 1999). Biopsy samples were taken from grey matter of the sensorimotor cortex and the grey matter of the thalamus, from one hemisphere for light microscopy and the contralateral for EM.

### Electron microscopy

Tissue for EM was post-fixed in a solution of 2% paraformaldehyde and 2.5% glutaraldehyde in 0.1M sodium cacodylate buffer. A version of the National Center for Microscopy and Imaging Research (NCMIR) protocol was adapted to provide enhanced contrast (Deerinck et al., 2010). Specifically, after fixation, the tissue underwent a multi-step enhanced contrast protocol at room temperature including sequential treatments with tannic acid, reduced osmium, thiocarbohydrazide, osmium, and uranyl acetate. This was followed by lead aspartate staining at 60°C. The brain sample was then dehydrated through a graded ethanol series, infiltrated with Embed 812 epoxy resin (EMS), and polymerized for 72 h at 60°C. Semithin sections (0.5 μm) were cut using a Leica UC7 ultramicrotome (Leica, Buffalo Grove, IL) and counterstained with 1% toluidine blue to identify the regions of interest within layers. Images were taken on a HT7500 transmission electron microscope (Hitachi High-Technologies, Tokyo, Japan) using an AMT NanoSprint12 12-megapixel CMOS TEM Camera System, software version 7.0.1.485 (Advanced Microscopy Techniques, Danvers, MA). Images were only adjusted for contrast on the AMT software. For three of the samples, we performed serial section transmission electron microscopy (ssTEM), using ultra-thin sections of 80 nm thickness, collected onto nickel slot grids. There were 6 image series from the identified region of interest across 10 serial sections.

We first performed fine-scale manual annotation of a subset of EM images, including one high-magnification image from all 20 unique brain samples (from different donors and brain regions). We annotated the boundaries of four mutually exclusive classes of structures across the image: (a) cell bodies, (b) membrane-bound structures with electron-dense interior, (c) membrane-bound structures with electron-lucent interior, and (d) myelinated axons. Category (c) is expected to include fluid-filled, swollen astrocyte processes, which is a common artifact seen in postmortem brain tissue, but is not necessarily expected to make it more difficult to trace neural processes (Krassner et al., 2023). The remainder of the image is designated as AIZs. These extracellular spaces may or may not contain partially degraded cellular membranes or other structural elements that cannot be reliably visualized due to postmortem changes or preparation artifacts. To assess inter-rater reliability for the fine-scale manual annotation, two independent raters annotated the same image. The spatial agreement between their annotations was quantified using the Dice similarity coefficient, with the median pairwise Dice score calculated across all polygons that were identified as having the most similar coordinates between the annotations.

We next graded two-dimensional EM images on whether or not they had AIZ artifacts, rating each image as either having or not having both (a) extensive AIZs and (b) a low degree of cell membrane intactness. We added the additional criterion of (b) to ensure that the images did not merely have expanded extracellular space, which could theoretically occur in conjunction with clearly delineated, well-preserved lipid membranes, for example depending upon the osmolarity of the preservative solutions (Pallotto et al., 2015). Two raters worked together to determine the grade for each image and resolved any discrepancies via consensus review. We calculated the percentage of images graded as having AIZ artifacts across each of the 5 medium-magnification EM images available for all 20 samples.

### Light microscopy

Brain tissue sampled for light microscopy was placed into cassettes for processing and embedded in paraffin. Paraffin-embedded brain sections 6 µm thick were baked, deparaffinized, and stained for H&E. Chromogenic immunohistochemistry was performed on Ventana Discovery Ultra according to the manufacturer’s directions (Neuropathology Brain Bank and Research CoRe, at Mount Sinai). The slides 6 µm thick were baked, deparaffinized, and pretreated using Cell Conditioning (CC1) antigen retrieval buffer (Tris/Borate/ EDTA buffer, pH 8.0–8.5, 950-224, Roche Diagnostics). Primary antibodies (**Table 1**) were diluted in antibody dilution buffer (ADB250, Ventana Medical System Inc., Roche Diagnostics). The detection was performed using the ultraView Universal DAB Detection Kit (760-500, Roche Diagnostics).

**Table 1.** List of antibodies used in this study.

| Antibody | Company | Catalog # | Dilution |
| --- | --- | --- | --- |
| Vimentin (V9) | Roche, Cell Marque | 05278139001 | 2.5ug/ml |
| GFAP (EP672Y) | Roche, Cell Marque | 5269784001 | 1ug/ml |
| NFL SMI-311 | BioLegend | 837904 | 1/1000 |
| NFL SMI-312 | BioLegend | 837801 | 1/1000 |

Hematoxylin and Bluing reagent (760-2021, 760-2037, Roche Diagnostics) were used as a nuclear counterstain. Digital images of the stained sections were captured at 40X as whole slide images (WSIs) using the Aperio GT450 high-resolution scanner (Leica Biosystems).

For the H&E- and immunohistochemically-stained WSIs, we first performed a qualitative assessment of preservation quality, examining the images for whether cell membrane morphology appeared intact across samples and also for common postmortem artifacts. For semi-quantitative grading of the samples, two raters collaboratively evaluated each sample and decided upon consensus grades. Perfusion quality on H&E staining was graded on a 0-3 scale by examining for the presence of intravascular material (including blood cells) across the WSI, with 0 indicating minimal intravascular material and 3 indicating extensive intravascular material. This analysis was done with the raters blinded to the preservation method used. GFAP immunostaining in each WSI was graded on a 0-3 scale based on staining intensity and quality, with 0 indicating poor staining (i.e., minimal or absent GFAP-positive structures) and 3 representing ideal staining patterns (i.e., widespread, clear visualization of astrocytic cell bodies and processes). We attempted to perform a similar grading scale for SMI-311 and SMI-312 immunostaining; however, we were unable to identify enough variability between the samples, as they all appeared to have robust and consistent immunoreactivity, with the exception of SMI- 311 for the two canine samples, which had negligible immunostaining.

## Results

### Preservation quality in electron microscopy images

Our cohort consisted of nine human brains and one canine brain, which we selected following their preservation to have a wide range of PMIs, from 1.5 hours to 4 days (**Table 2**). For each brain, biopsy samples were obtained from the grey matter of the sensorimotor cortex and thalamus.

**Table 2.** Characteristics of brain donors included in this study. Fixation duration refers to the amount of time in 10% neutral buffered formalin at 4°C. *: This donor utilized medical aid in dying (MAID) for end-of-life care. **: Canine brain. PMI: Postmortem interval.

| Donor ID | Age | Sex | PMI | Reported cause of death | Preservation method | Duration of fixation |
| --- | --- | --- | --- | --- | --- | --- |
| 7 | 78 | Male | 4.25 hours | Cancer* | Perfusion fixation | 5 months |
| 30 | 77 | Male | 17 hours | Kidney failure, failure to thrive | Perfusion fixation | 4 months |
| 34 | 88 | Male | 20 hours | Cancer* | Perfusion fixation | 4 months |
| 37 | 70 | Male | 72 hours | Myocardial infarction | Perfusion fixation | 4 months |
| 55 | 70 | Male | 36 hours | Cardiac causes | Perfusion fixation | 3 months |
| 57 | 75 | Male | 96 hours | Leukemia | Immersion fixation | 3 months |
| 59 | 41 | Female | 91 hours | Leukemia | Perfusion fixation | 2.5 months |
| 65 | 17** | Female | 1.5 hours | Euthanasia (Donated canine) | Perfusion fixation | 2 months |
| 66 | 89 | Female | 61 hours | Sepsis and acute renal failure | Perfusion fixation | 2 months |
| 78 | 89 | Male | 2.5 hours | Failure to thrive | Perfusion fixation | 1 month |

Qualitatively, we observed varying degrees of ultrastructural preservation across the specimens. Across all samples, we detected several artifacts commonly found in postmortem brain tissue, including cellular swelling and shrinkage, vacuolization, myelin disbanding, and partially disrupted cell membranes whose potential shapes could still be at least partially distinguished (Krassner et al., 2023). The canine brain, which had the shortest PMI of 1.5 hours, was found to have the lowest burden of these postmortem artifacts.

We next performed fine-scale manual annotation of ultrastructural components in the images (**Figure 1**). To assess the reproducibility of our annotation approach, two independent raters annotated the same image, which was from the cortex sample of donor #66. Inter-rater reliability was quantified by calculating the Dice similarity coefficient for each pair of matching polygons, yielding a median Dice score of 0.79 (**Figure 2**). In one high-magnification image from each unique sample, we also quantified the percentage of area occupied by AIZs (**Figure 3**, **Table 3**).

**Figure 1.**
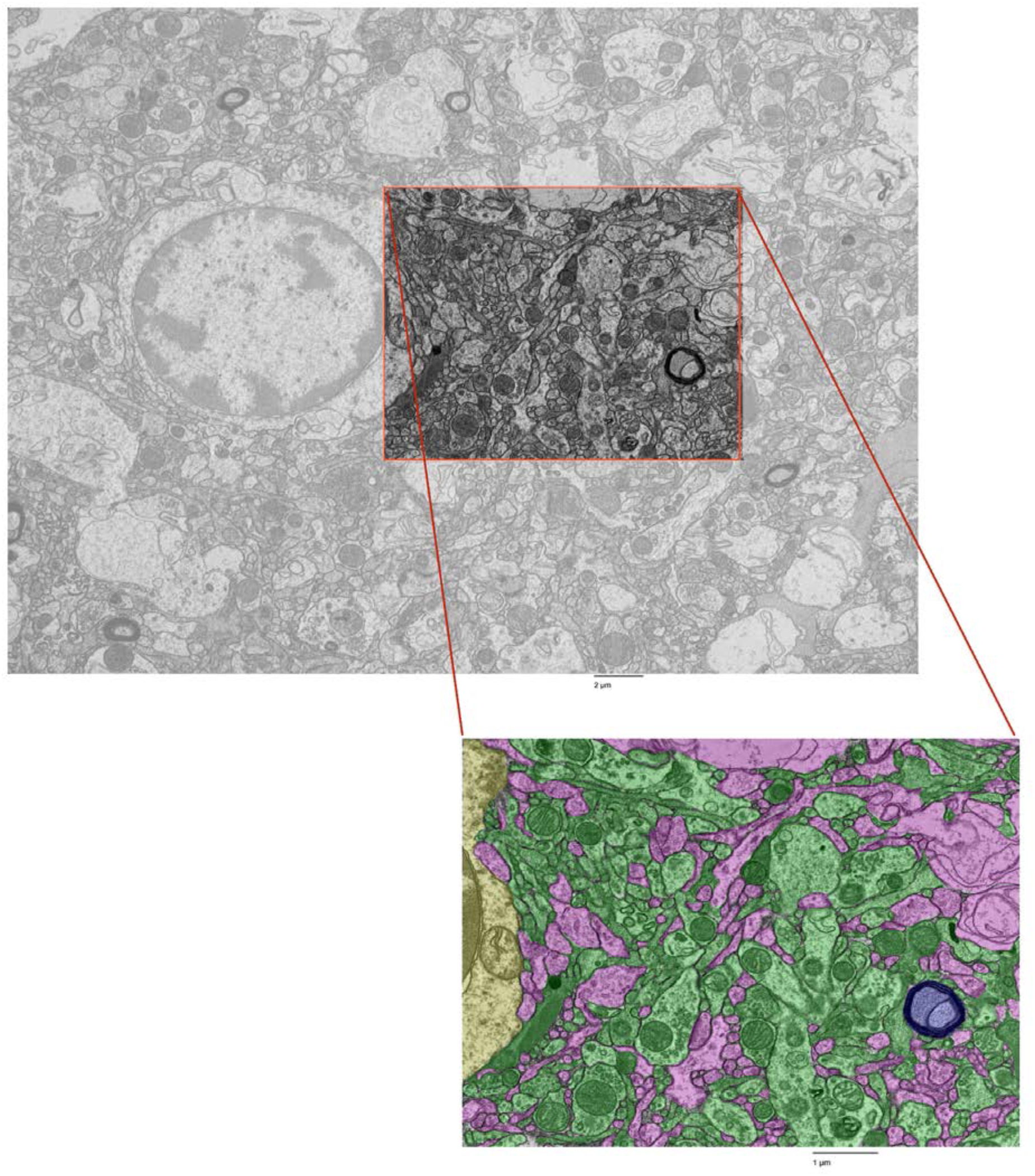
Fine-scale annotation of ultrastructural components in one representative EM image from the cortex of donor #7. Color coding identifies distinct cellular structures: Green: membrane-bound structures with electron-dense interior (e.g., organelle-rich neurites); Purple: membrane-bound structures with electron-lucent interior (e.g., swollen astrocytic processes); Blue: myelinated axons; Yellow: cell bodies. Non-colored regions represent ambiguous interstitial zones (AIZs) that lack clearly defined membrane boundaries. Upper scale bar: 2 µm. Lower scale bar: 1 µm.

**Figure 2.**
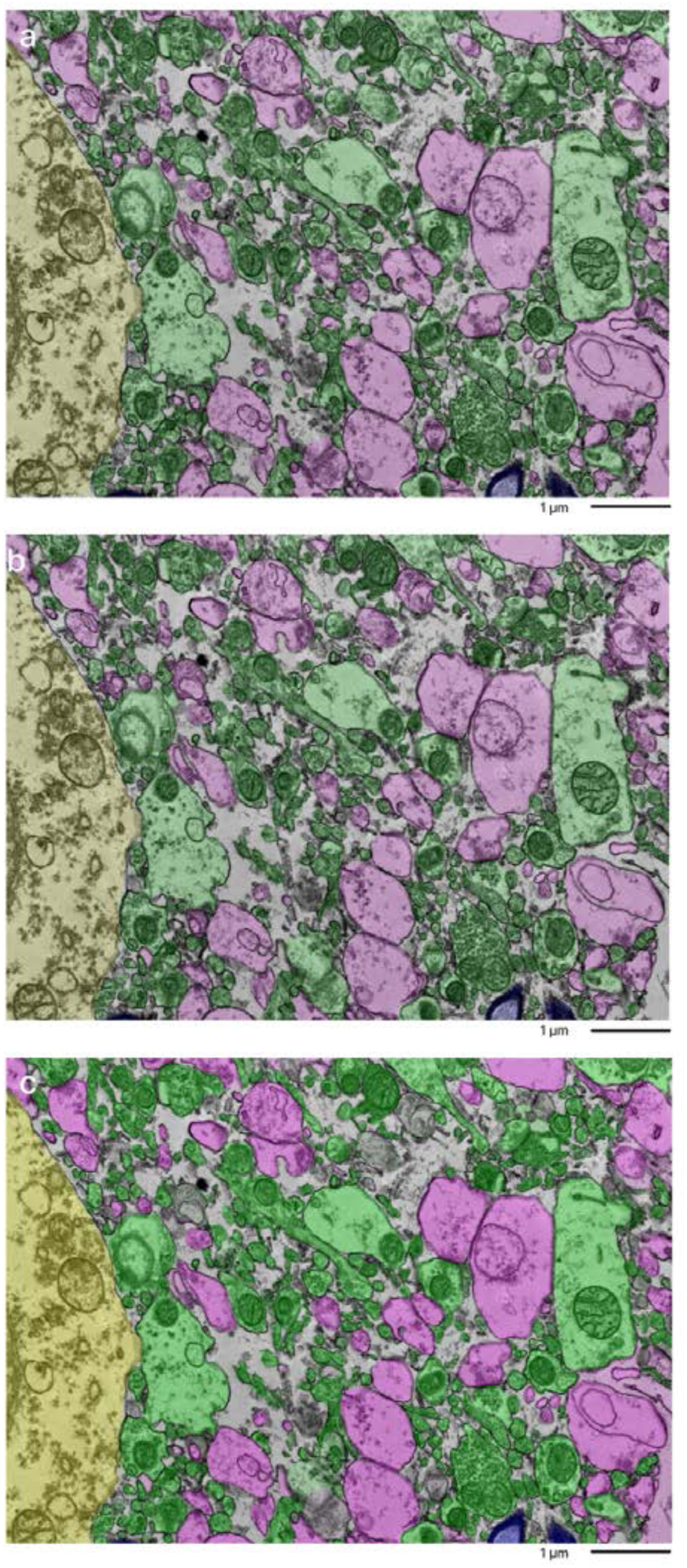
Overlap of annotations of a sample from the cortex of donor #66 by independent annotators. The top and middle image were done by two independent annotators. The bottom image is a combination of the overlays to compare how alike the two annotations are. Color coding identifies distinct cellular structures: Green: membrane-bound structures with electron-dense interior (e.g., organelle-rich neurites); Purple: membrane-bound structures with electron-lucent interior (e.g., swollen astrocytic processes); Blue: myelinated axons; Yellow: cell bodies. Non-colored regions represent ambiguous interstitial zones (AIZs) that lack clearly defined membrane boundaries. Scale bars: 1 µm.

**Figure 3.**
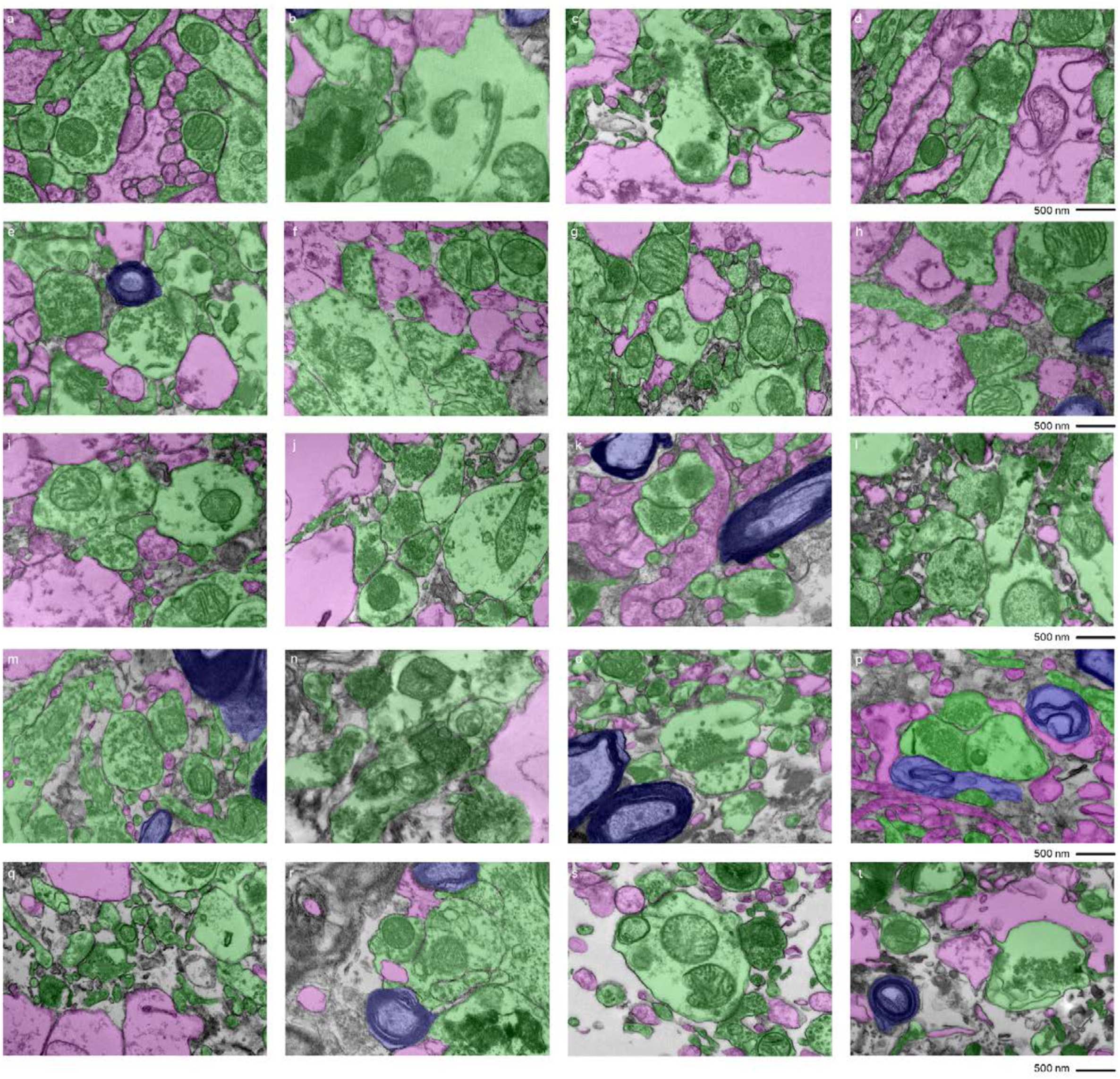
Fine-scale manual annotation of ambiguous interstitial zones (AIZs) from one EM image across all samples. Samples are sorted by the percentage of ambiguous interstitial zones (AIZs), from the lowest to the highest: 7-C (**a**), 37-T (**b**), 34-C (**c**), 65-C (**d**), 59-C (**e**), 55-C (**f**), 37-C (**g**), 78-C (**h**), 57-C (**i**), 30-C (**j**), 65-T (**k**), 57-T (**l**), 55-T (**m**), 78-T (**n**), 30-T (**o**), 34-T (**p**), 66-C (**q**), 7-T (**r**), 66-T (**s**), 59-T (**t**), where the number is the donor ID number, and then “-T” indicates that it is from the thalamus and “-C” from the cortex. Color coding identifies distinct cellular structures: Green: membrane-bound structures with electron-dense interior (e.g., organelle-rich neurites); Purple: membrane-bound structures with electron-lucent interior (e.g., swollen astrocytic processes); Blue: myelinated axons; Yellow: cell bodies. Non-colored regions AIZs that lack clearly defined membrane boundaries. Scale bars: 1 µm.

**Table 3.**
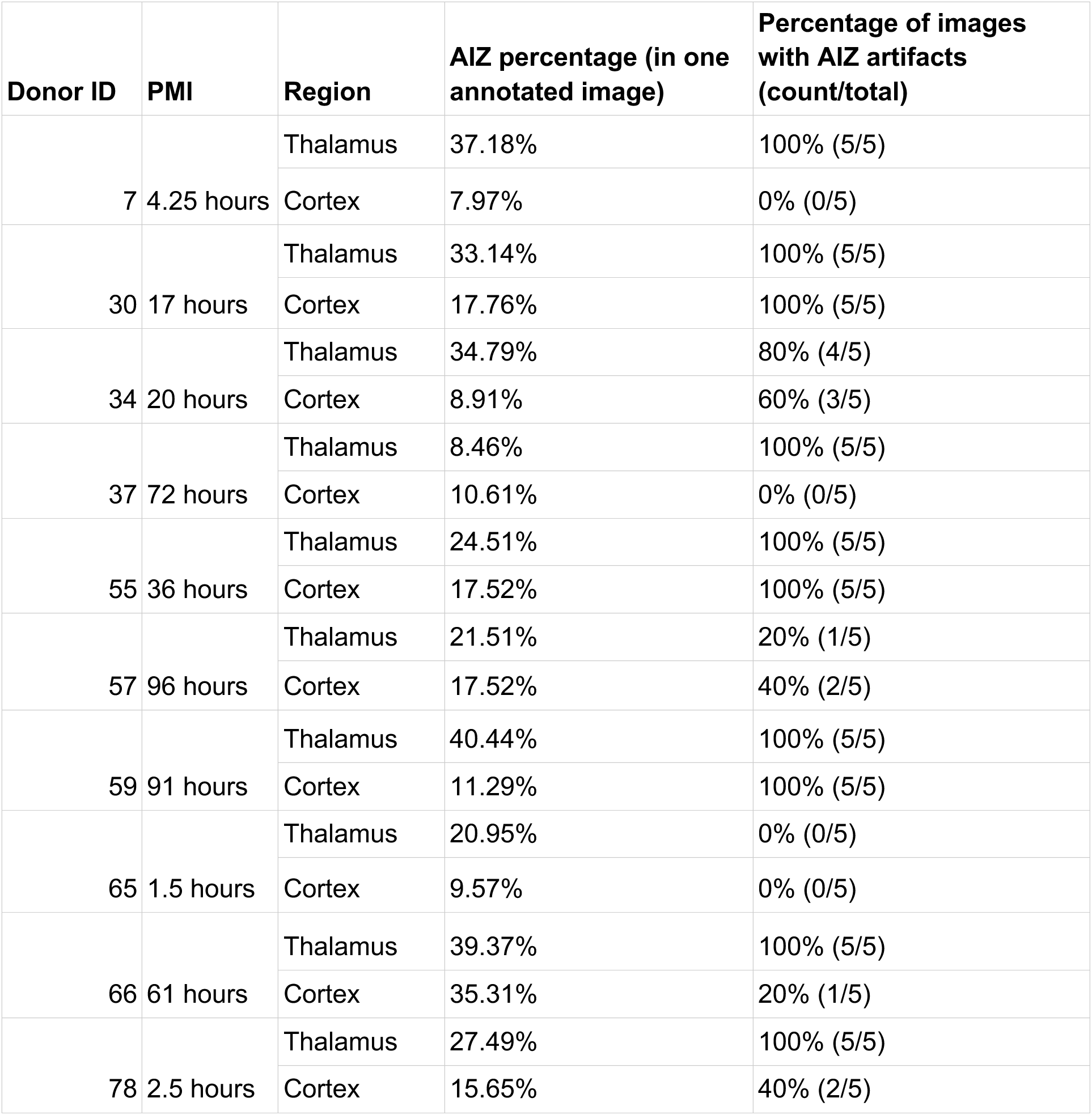
Summary of tissue preservation quality metrics by donor and brain region. The “AIZ percentage” column shows the proportion of area occupied by AIZs in one high magnification, annotated EM image, representing electron-lucent, non-membrane-bound regions that may contain degraded cellular structures. The “Percentage of images with AIZ artifacts” column indicates how many EM images were graded as having AIZ artifacts, with the raw number of affected images shown in parentheses (out of 5 total images evaluated per sample). PMI: Postmortem interval; AIZ: Ambiguous interstitial zone.

| Donor ID | PMI | Region | AIZ percentage (in one annotated image) | Percentage of images with AIZ artifacts (count/total) |
| --- | --- | --- | --- | --- |
| 7 | 4.25 hours | Thalamus | 37.18% | 100% (5/5) |
|  |  | Cortex | 7.97% | 0% (0/5) |
| 30 | 17 hours | Thalamus | 33.14% | 100% (5/5) |
|  |  | Cortex | 17.76% | 100% (5/5) |
| 34 | 20 hours | Thalamus | 34.79% | 80% (4/5) |
|  |  | Cortex | 8.91% | 60% (3/5) |
| 37 | 72 hours | Thalamus | 8.46% | 100% (5/5) |
|  |  | Cortex | 10.61% | 0% (0/5) |
| 55 | 36 hours | Thalamus | 24.51% | 100% (5/5) |
|  |  | Cortex | 17.52% | 100% (5/5) |
| 57 | 96 hours | Thalamus | 21.51% | 20% (1/5) |
|  |  | Cortex | 17.52% | 40% (2/5) |
| 59 | 91 hours | Thalamus | 40.44% | 100% (5/5) |
|  |  | Cortex | 11.29% | 100% (5/5) |
| 65 | 1.5 hours | Thalamus | 20.95% | 0% (0/5) |
|  |  | Cortex | 9.57% | 0% (0/5) |
| 66 | 61 hours | Thalamus | 39.37% | 100% (5/5) |
|  |  | Cortex | 35.31% | 20% (1/5) |
| 78 | 2.5 hours | Thalamus | 27.49% | 100% (5/5) |
|  |  | Cortex | 15.65% | 40% (2/5) |

We next developed a grading method to rate EM images for AIZs that could both be performed more rapidly by trained annotators and could also take into account the degree of cell membrane intactness. Specifically, the images were graded as having AIZ artifacts if they displayed both (a) extensive AIZs and (b) poor delineation of cellular membrane boundaries (**Figure 4**). This combined assessment is meant to distinguish artifactual AIZs from potential scenarios with non-artifactual extracellular space expansion. The frequency of images meeting these criteria for having AIZ artifacts was calculated for each sample (**Table 3**). We found that samples from a donated canine preserved after a PMI of 1.5 hours yielded EM images with no AIZ artifacts in either brain region. The brains from two of the human donors, one with a PMI of 4.5 hours (donor #7) and one with a PMI of 72 hours (donor #37), also yielded EM images with no AIZ artifacts in the cortical samples, but both of these brains had AIZ artifacts detected in images from the thalamus samples. The cortex and thalamus samples from the other 7 brains produced EM images graded as having AIZ artifacts in at least one of the five available EM images.

**Figure 4.**
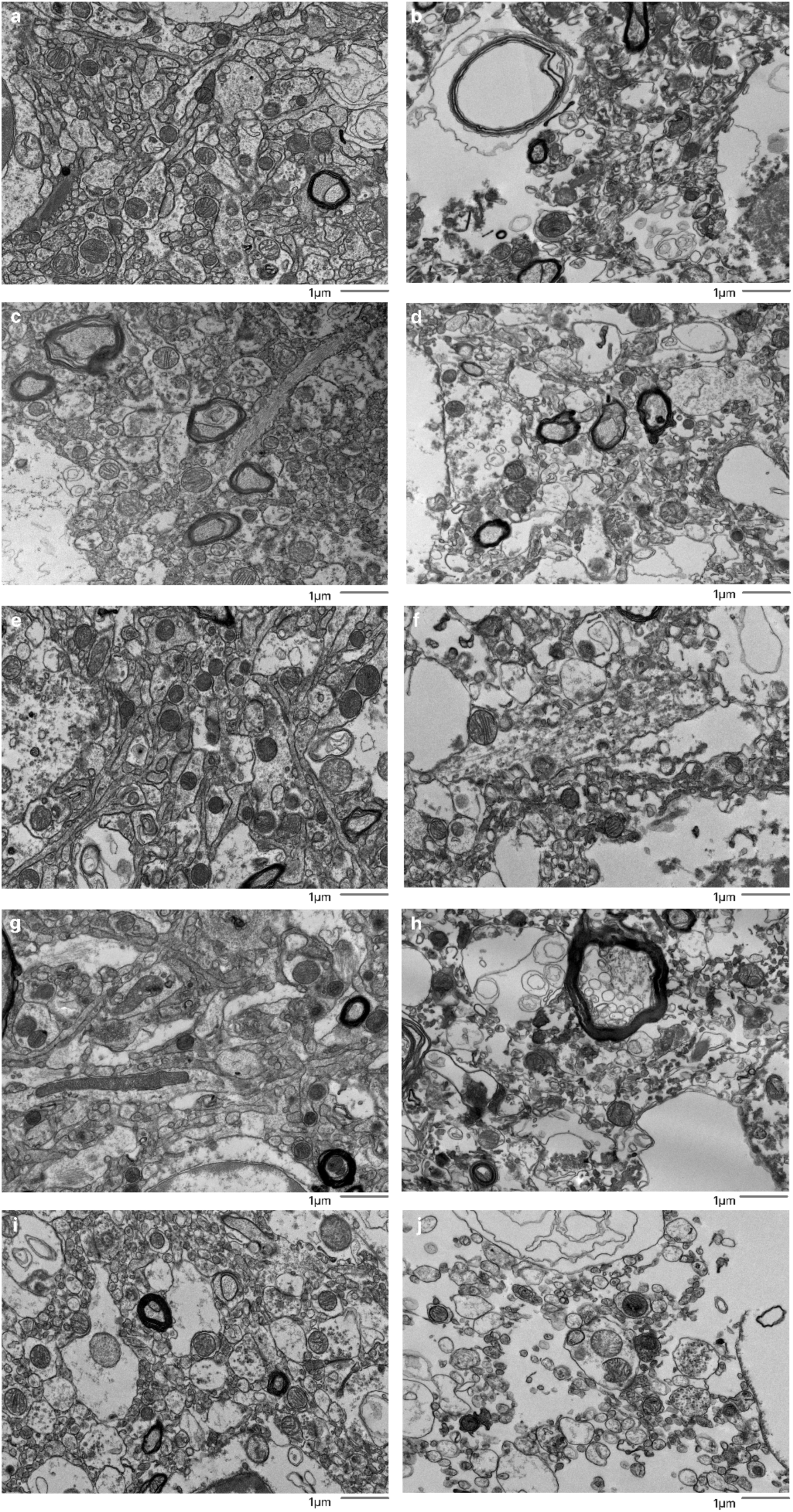
Representative EM images graded as having or not having of AIZ artifacts. Five examples images are shown that were graded as having AIZ artifacts: 30-T (**b**), 39-C (**d**), 59-C (**f**), 59-T (**h**), 66-T (**j**), as well as five examples that were not: 7-C (**a**), 57-C (**c**), 65-C (**e**), 65-T (**g**), 66-C (**i**), where the number is the donor ID number, and then “-T” indicates that it is from the thalamus and “-C” from the cortex. Scale bars: 1 µm. AIZ: Ambiguous interstitial zone.

We found that there was no significant rank correlation between the PMI and the annotated AIZ percentage in the images from either the cortex (rho = 0.35, p = 0.327) or the thalamus (rho = 0.09, p = 0.811). Similarly, we found no significant rank correlation between the PMI and the percentage of images graded as having AIZ artifacts in the samples from either the cortex (rho = 0.27, p = 0.454) or the thalamus (rho = 0.10, p = 0.790). Next, we found that there was a significantly higher annotated AIZ percentage in images from the thalamus than the cortex (mean in thalamus = 28.8%, mean in cortex = 15.2%, t-test p-value = 0.004). This result may reflect differences in preservation quality, baseline differences between the regions, or other factors. There was also a non-significant regional trend in the percentage of images graded as having AIZ artifacts, which was higher in the thalamus than the cortex (mean in thalamus = 80%, mean in cortex = 46%, t-test p-value = 0.074).

We next performed a qualitative analysis of the ssTEM data, which was derived from cortical samples. First, synapses were identified in the available image stacks, all of which could be traced through the image stack until synapse termination or the end of image stack, without apparent loss of structure (**Figure 5**). Next, we attempted to trace the neural connections arising from synapses to their associated dendrites and axons across the image stacks. Qualitatively, we found that for the sample from donor #35, which had a more extensive burden of AIZ artifacts on the 2D EM images, it was more difficult to manually trace the neurite structures. For the other two samples imaged with ssTEM (donor IDs #7 and #65), we successfully identified traceable neurites across the image stack in at least some instances (for example, **Figure 6**).

**Figure 5.**
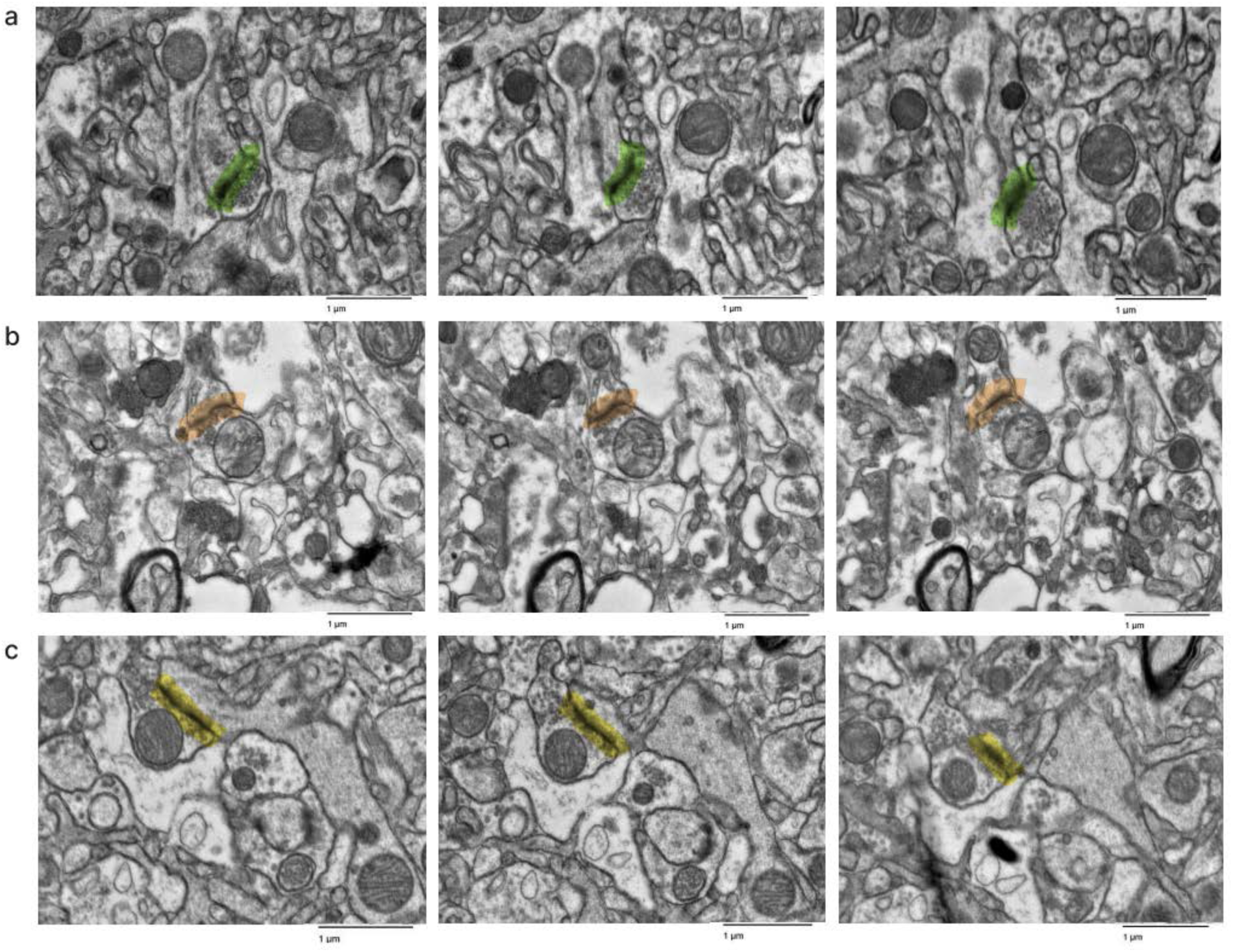
Representative images in the serial section TEM data sets showing that synapses can be traced. Samples from the cortex of donor IDs #7 (**a**), #35 (**b**), #65 (**c**). Synapses are highlighted in color. Scale bars: 1 µm.

**Figure 6.**
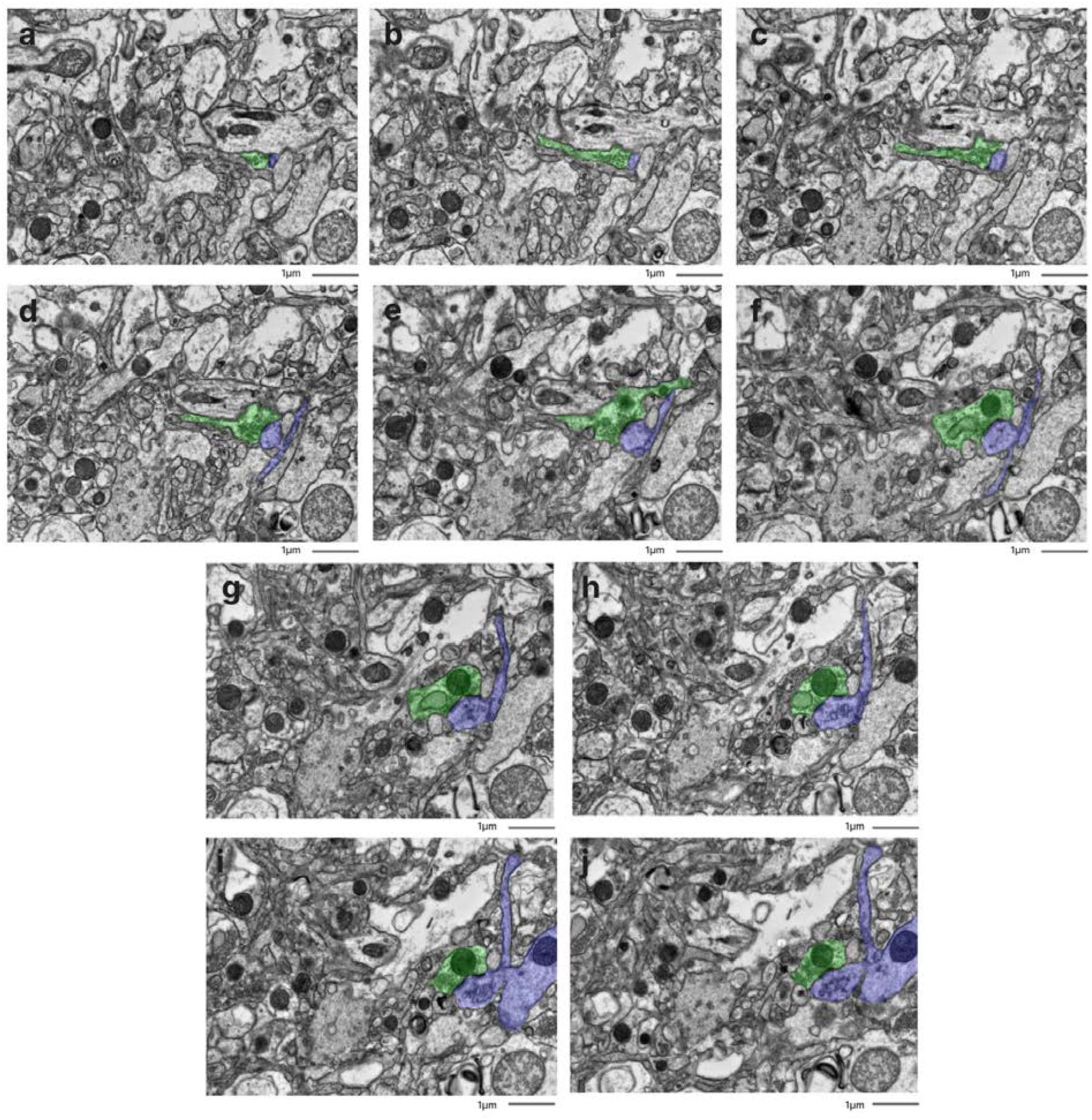
Representative serial section image stack with annotations for afferent neurites from one synapse. All images are from a serial section from the cortex sample of donor #65. Scale bars: 1 µm.

However, we were unable to rigorously quantify the traceability due to two key limitations: (a) an insufficient sample size to establish reliable metrics, and (b) technical constraints related to our section thickness. Specifically, the resolution between sequential images made it challenging to differentiate between potential structural damage and normal morphological variations where processes might naturally terminate or significantly change direction.

### Preservation quality in light microscopy images

We next set out to compare preservation quality between modalities. We started by performing an analysis of H&E-stained light microscopy sections. These samples were obtained from the contralateral hemisphere of the same brain regions that were examined by electron microscopy. We detected expected postmortem changes in all images, including pericellular rarefactions, perivascular rarefactions, and neuropil vacuolization (Krassner et al., 2023). Qualitatively, the extent of pericellular rarefaction and neuropil vacuolization appeared to potentially be more pronounced in some cases with longer PMIs. However, no obvious differences in the preservation quality of cell membrane morphology were observed that corresponded with the PMI on these H&E-stained images (**Figure 7**). Taken together, these findings suggest that basic cellular morphology remains largely intact at the light microscopy level in the samples from these brains, despite the range of PMIs.

**Figure 7.**
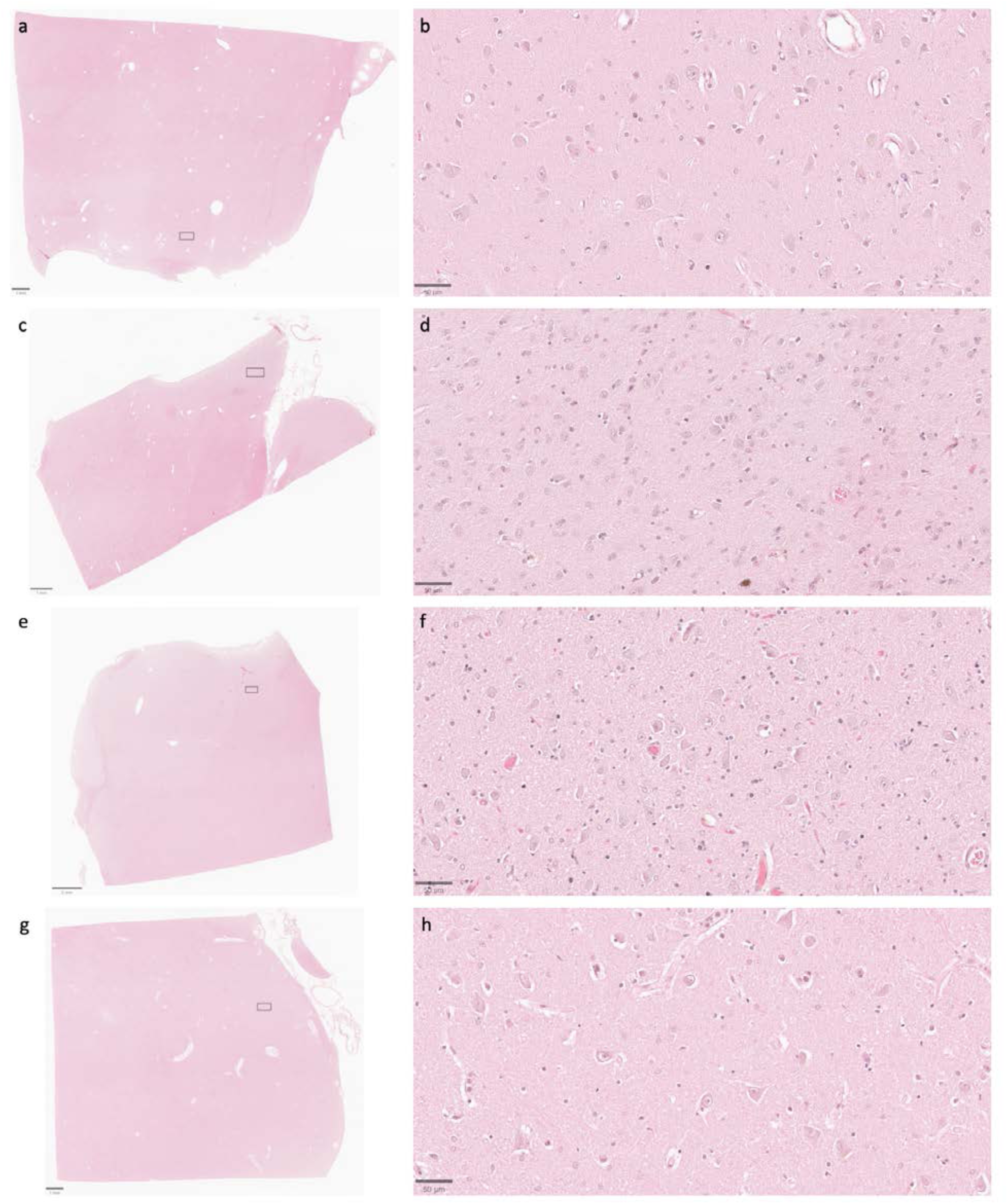
Representative H&E-stained images demonstrate that cell membrane morphology is generally intact across a wide range of PMIs. Samples from the thalamus of donor IDs #7 (**a**, **b**), #65 (**c**, **d**), #57 (**e**, **f**), and #59 (**g**, **h**) with PMIs of 4.5 h, 1.5 h, 96 h, and 91 h, respectively.

Insets on the lower magnification images correspond to the regions shown in the higher magnification images. Scale bars in (**a**, **c**, and **g**) are 1 mm; scale bar in **e** is 2 mm; scale bars in (**b**, **d**, **f**, and **h**) are 50 μm.

Notably, in cases where perfusion fixation was performed, the extent to which perfusate was successfully delivered to the brain regions examined varied across cases, as evidenced by differences in brain stiffness and the extent of blood clearance from surface vessels. This variability is consistent with previous findings on postmortem brain perfusion (McFadden et al., 2019). As one additional way to measure this, we assessed H&E-stained images to measure the degree to which blood vessels were cleared of material such as red blood cells (**Figure 8**, **Table 4**). These data demonstrate that blood vessels were not fully cleared, and therefore perfusion was not complete, in the majority of the brains. This is further supported by the observation that the immersion-fixed brain (donor #57) had a similar grade for intravascular material clearance as several of the perfusion-fixed samples. Taken together, these results suggest that in cases where a perfusion procedure was performed, the fixative may not have fully penetrated the vasculature throughout the brain, perhaps instead only traversing through a subset of blood vessels.

**Figure 8.**
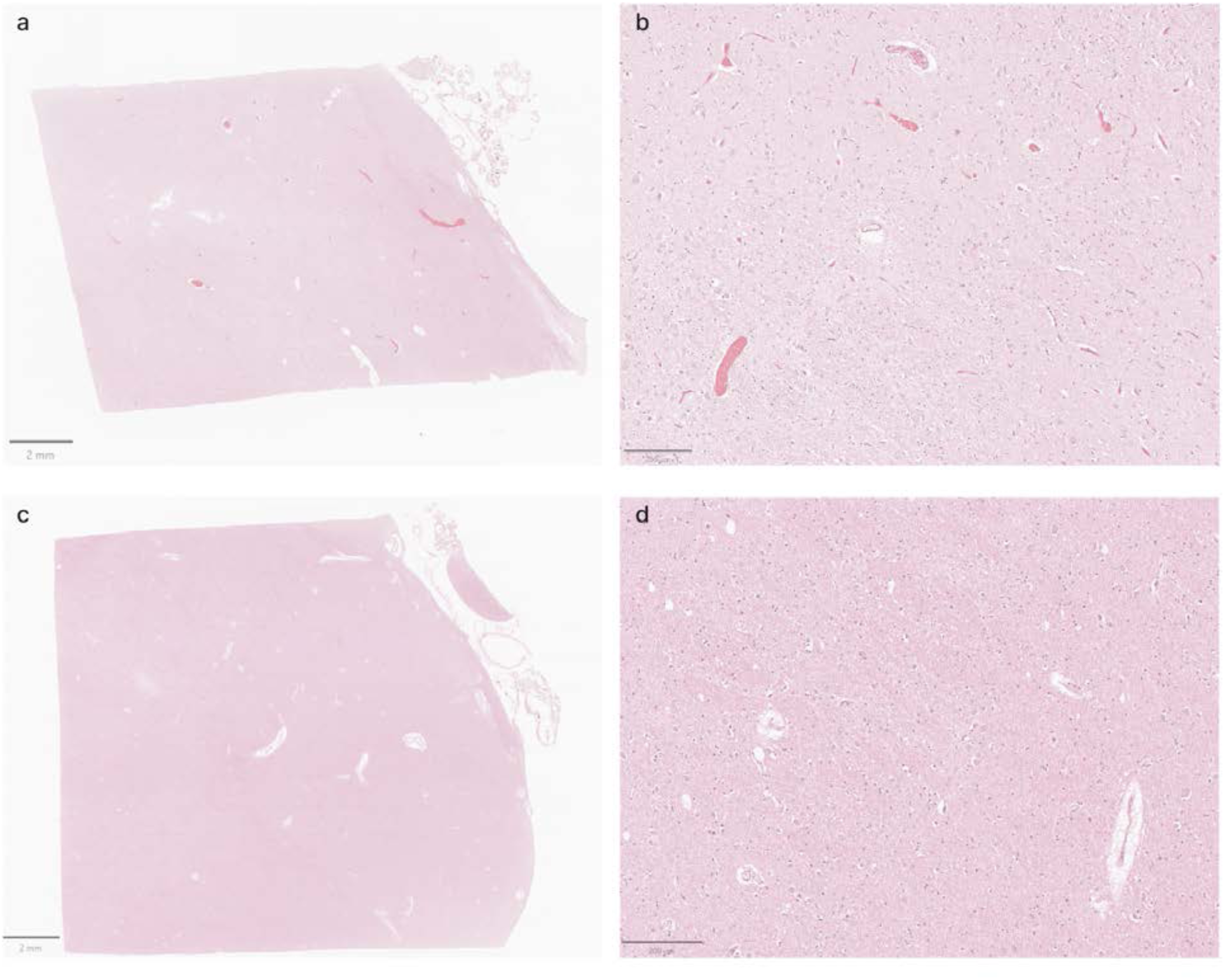
Representative H&E-stained images from the thalamus showing variable vascular clearance. **a**, **b**: Images from donor #30 shows extensive intravascular material. **c**, **d**: Images from donor #59 shows minimal intravascular material. Scale bars: (**a**, **c**) 2 mm; (**b**, **d**) 200 μm.

**Table 4.** Semi-quantitative grades to evaluate preservation quality on light microscopy. The presence of intravascular material was graded on a scale of 0-3, where 0 indicates minimal intravascular material and 3 indicates extensive intravascular material. GFAP staining was graded on a scale of 0-3, where 0 indicates minimal or absent GFAP-positive structures and 3 represents widespread visualization of astrocytic cells and processes.

| Donor ID | PMI | Region | Intravascular material grade on H&E images | GFAP staining grade |
| --- | --- | --- | --- | --- |
| 7 | 4.25 hours | Thalamus | 1 | 2 |
|  |  | Cortex | 1 | 3 |
| 30 | 17 hours | Thalamus | 3 | 1 |
|  |  | Cortex | 1 | 0 |
| 34 | 20 hours | Thalamus | 2 | 1 |
|  |  | Cortex | 2 | 1 |
| 37 | 72 hours | Thalamus | 2 | 0 |
|  |  | Cortex | 2 | 0 |
| 55 | 36 hours | Thalamus | 2 | 0 |
|  |  | Cortex | 1 | 2 |
| 57 | 96 hours | Thalamus | 2 | 1 |
|  |  | Cortex | 2 | 3 |
| 59 | 91 hours | Thalamus | 0 | 0 |
|  |  | Cortex | 2 | 0 |
| 65 | 1.5 hours | Thalamus | 2 | 3 |
|  |  | Cortex | 1 | 3 |
| 66 | 61 hours | Thalamus | 2 | 0 |
|  |  | Cortex | 2 | 1 |
| 78 | 2.5 hours | Thalamus | 1 | 2 |
|  |  | Cortex | 1 | 3 |

We next performed immunohistochemical staining to evaluate the preservation of specific neural components across our samples. Staining for SMI-312, a pan-axonal stain targeting highly phosphorylated neurofilaments, revealed consistent preservation of axonal architecture across samples with varying PMIs (**Figure 9**) (Ulfig et al., 1998). Even in samples with PMIs exceeding 60 hours, we observed well-defined axonal processes with clear continuity and distinct morphology. In the cortical samples, neurofilament-positive axons were readily identifiable traversing throughout the neuropil, with minimal disruption of their trajectory detected on qualitative analysis.

**Figure 9.**
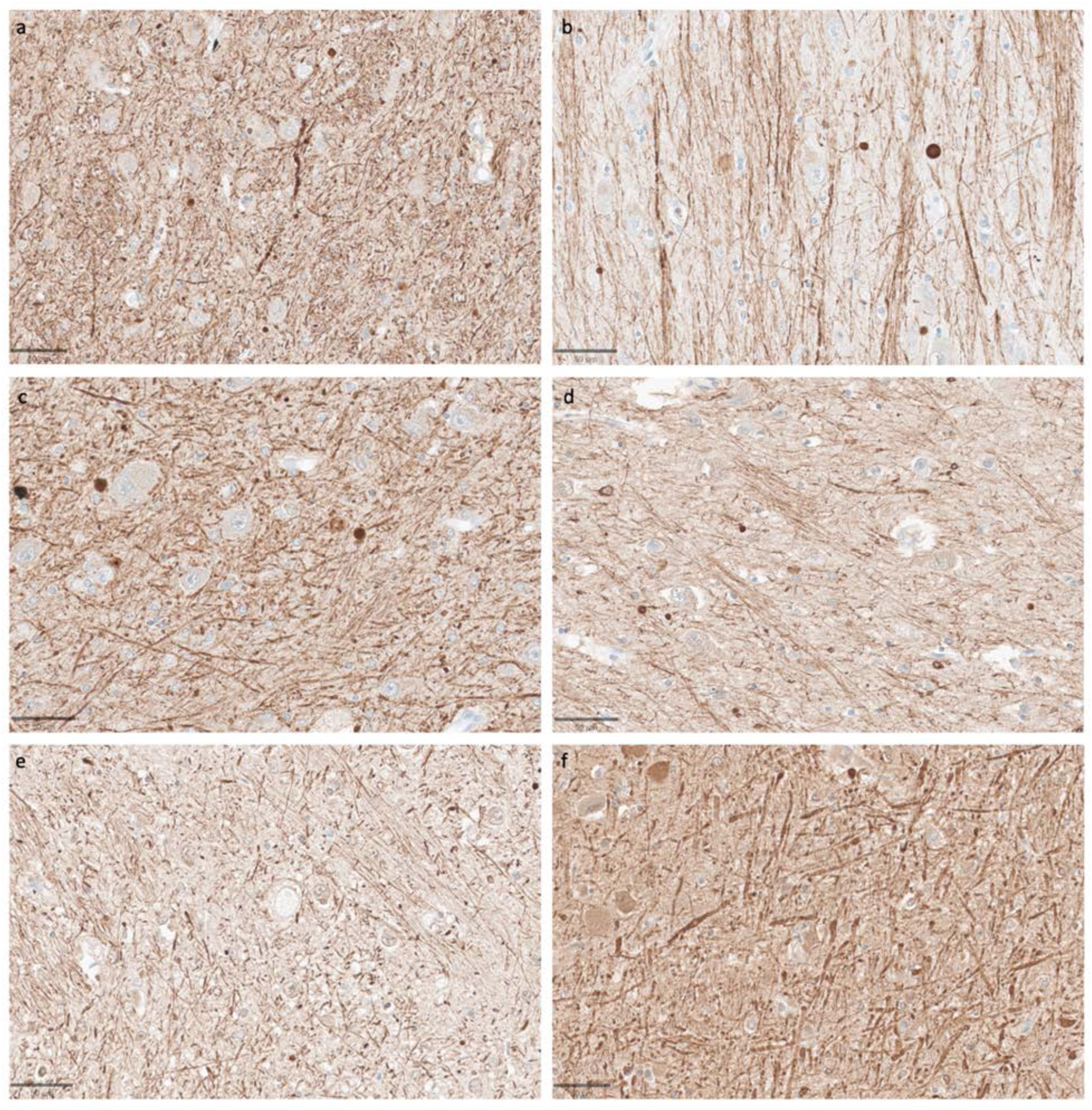
Representative SMI-312-stained images demonstrate that axonal morphology is generally intact across a wide range of PMIs. Samples from the thalamus of donor IDs #65 (**a**), #78 (**b**), #7 (**c**), #37 (**d**), #59 (**e**), and #57 (**f**) with PMIs of 1.5 h, 2.5 h, 4.5 h, 72 h, 91 h, and 96 h, respectively. Scale bars: 50 μm.

Staining for SMI-311, an antibody targeting non-phosphorylated neurofilaments that are primarily located in the dendrites and perikarya, also revealed consistent staining quality across samples with varying PMIs (**Figure 10**) (Ulfig et al., 1998). Dendritic morphology appeared to be largely intact, without the diffuse staining that would be expected with widespread breakdown of the dendritic cytoskeleton. A key exception was the canine brain, which did not stain adequately for this antibody in either brain region, possibly reflecting molecular differences in neurofilaments between human and canine brains (Dimakopoulos and Mayer, 2002).

**Figure 10.**
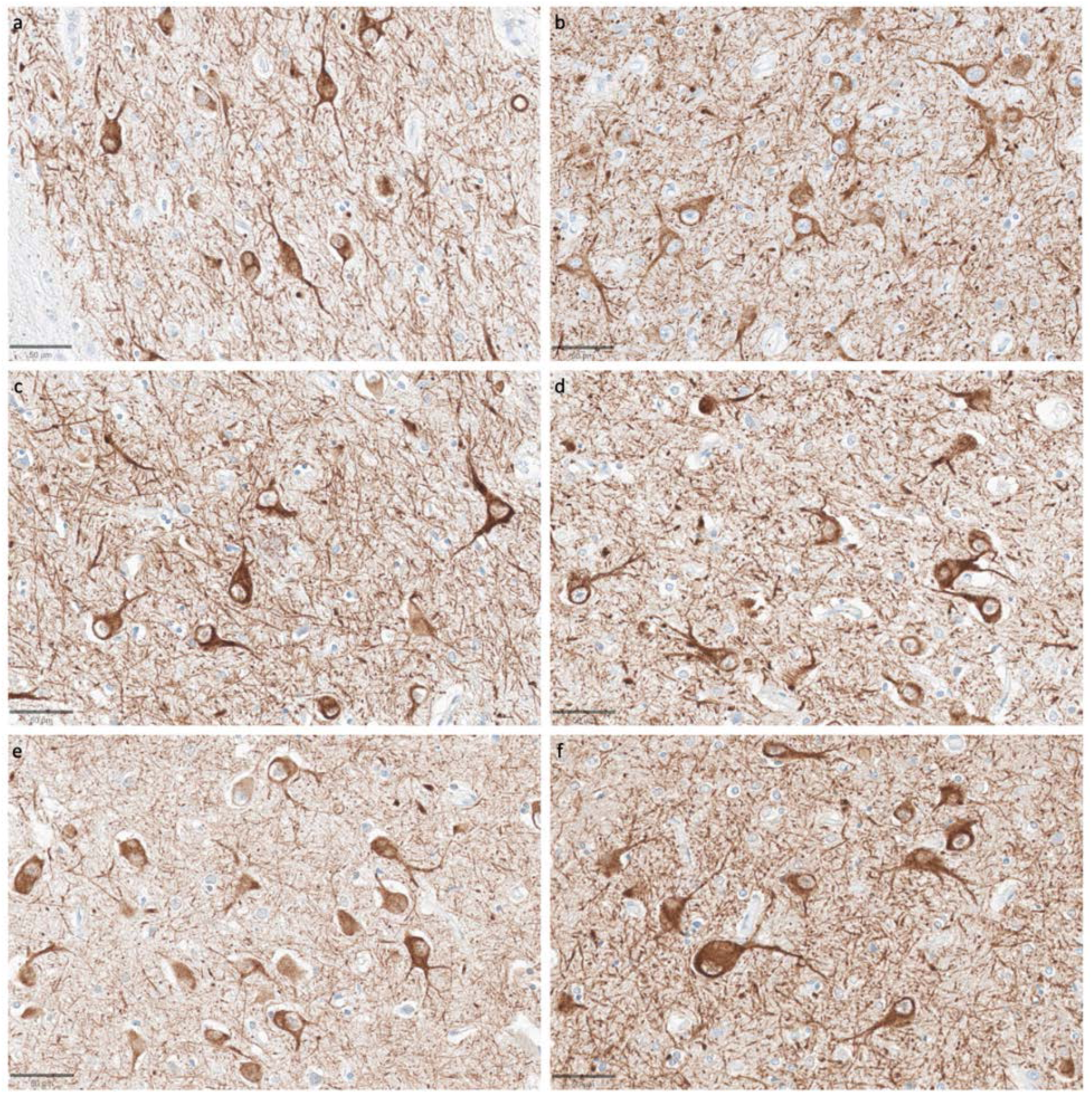
Representative SMI-311-stained images demonstrate that dendritosomatic morphology on light microscopy is generally intact across a wide range of PMIs in human brains. Samples from the thalamus of donor IDs #78 (**a**), #7 (**b**), #30 (**c**), #37 (**d**), #59 (**e**), and #57 (**f**) with PMIs of 2.5 h, 4.5 h, 17 h, 72 h, 91 h, and 96 h, respectively. Scale bars: 50 μm.

We next assessed the preservation of astrocyte morphology via immunostaining for the astrocyte marker GFAP. In samples from the thalamus, we observed that GFAP staining was generally lower compared to the cortical samples, with the exception of the subependymal area, which consistently showed higher immunoreactivity. In the cortical samples, the subpial area consistently had the highest density of staining, which is expected given previous findings about GFAP staining patterns (Halliday et al., 1996). The density of GFAP staining in the cortex appeared to be lower in some cases with longer PMIs, but there was no robust trend for lower GFAP staining with longer PMIs, as evidenced by strong staining in some samples that had relatively longer PMIs, such as one brain with a PMI of 96 hours (**Figure 11**). Moreover, when GFAP-positive astrocytes were visible in the longer PMI cases, the individual cells that were stained appeared to still have intact morphology without obvious signs of partial decomposition, arguing against cellular decomposition playing the major role in mediating staining differences. However, given our relatively small sample size, we cannot clearly distinguish the reasons for variability in GFAP staining across samples.

**Figure 11.**
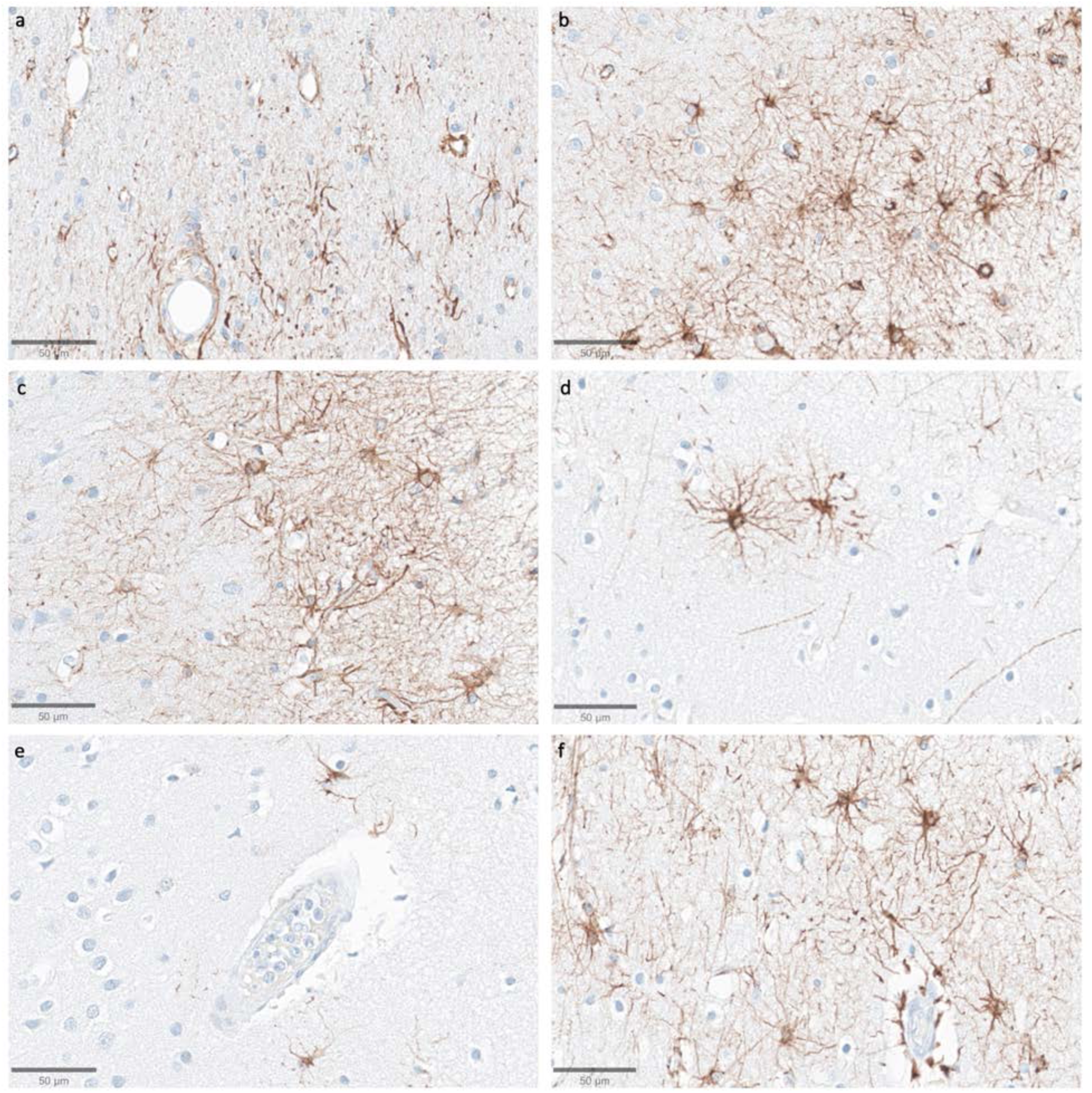
Representative GFAP-stained images of the cortex show intact subpial astrocyte morphology in images from samples with a wide range of PMIs. Samples from donor IDs #65 (**a**), #78 (**b**), #7 (**c**), #37 (**d**), #59 (**e**), and #57 (**f**) with PMIs of 1.5 h, 2.5 h, 4.5 h, 72 h, 91 h, and 96 h, respectively. Scale bars: 50 μm.

In a subset (n=6) of the samples, we also performed immunostaining with vimentin (see Data Availability for access to these WSIs). While this marker did stain around blood vessels, its staining for astrocytes appeared highly sensitive to the duration of fixation, more so than GFAP, showing only weak and inconsistent staining in samples fixed for longer than one month. Due to this limitation of vimentin staining, we did not extend the vimentin staining to the full cohort.

## Discussion

In this study, we first present a framework for evaluating preservation quality in postmortem brain tissue through the assessment of ultrastructural integrity in EM images. We found that AIZ artifacts occurred in at least one of the available EM images from our tested samples, with the exception of a canine sample with a relatively shorter PMI of 1.5 hours and two cortical samples from human donors. Notably, in the cortical sample from the canine and one of these two human cortical samples, we were able to trace the neurites arising from synapses across ssTEM image stacks in some instances, though technical limitations and insufficient sample volumes prevented rigorous quantification. Our observations suggest that samples without AIZ artifacts on 2D images may be a proxy for the degree of ultrastructural integrity that could potentially support connectomics studies, though this requires more imaging and larger sample sizes for corroboration. We also detected a trend towards a higher percentage of AIZs in thalamic samples compared to cortical ones, which may be statistical noise, may be attributable to a lower average preservation quality of the thalamus than the cortex in this cohort, or may reflect baseline regional anatomical differences that we have not fully accounted for. Next, we performed light microscopy on samples from the same brains, using both non-specific morphological staining as well as immunostaining for neurofilaments and GFAP. The subjective quality of the morphological staining and neurofilament staining appeared similar across samples from all 10 brains, despite the varying PMIs and differences observed on EM images. Our data suggests a complex relationship between the imaging findings and the underlying tissue preservation depending on the imaging modality used, which warrants further investigation.

There is an extensive literature on the visualization of extracellular space in brain tissue, with many aspects of the preservation procedure including the delivery technique, fixative, buffer, and osmolarity all significantly affecting it (Van Harreveld and Malhotra, 1967; Cragg, 1980; Fix and Garman, 2000; Korogod et al., 2015; Fulton and Briggman, 2021). Living brain tissue generally contains around 20% extracellular space (Nicholson and Syková, 1998). Standard methods of perfusion fixation in laboratory animals, which involves a minimal duration of ischemia prior to preservation, can decrease the extracellular space in an artifactual manner from its typical *in vivo* volume (Pallotto et al., 2015). Our findings suggest that although there may be an initial decrease in the ECS from *in vivo* levels with minimal PMI, which may involve ischemic cell swelling, as the PMI progresses, the ECS may once again grow to *in vivo* levels and beyond as fluid redistributes. Indeed, one would expect that given a sufficiently long PMI, the brain would be entirely “extracellular,” as all of the cellular structures would eventually break down and the brain liquefy. Although we expect that molecular decomposition and fluid redistribution is the most likely explanation for AIZs, there are several non-mutually exclusive explanations for the AIZs we observed in our samples with longer PMIs. Some degree of extracellular space is expected even in ideally preserved tissue, and may in fact be desirable as this better reflects *in vivo* conditions and can aid in neurite tracing (Korogod et al., 2015; Pallotto et al., 2015). However, the appearance of potential debris alongside the expansion of the extracellular space suggests a significant change from normal tissue architecture. This is why our grading method for AIZ artifacts takes into account both the presence of AIZs and a qualitative determination of poor cell membrane intactness.

One notable pattern we observed in our data is that the occurrence of AIZs artifacts qualitatively appears to coincide with other postmortem changes involving fluid shifts. For example, images with AIZs artifacts also tend to exhibit increased astrocyte process swelling, vacuolization, and cytoplasmic washout. This pattern is evident in both our images and in other publicly available electron microscopy data sets of postmortem human brain tissue (Oost et al., 2023). These observations suggest that there may be a generalized “excess fluid” phenomenon in some postmortem brain tissue, with AIZs artifacts being just one manifestation of this. Theoretically, one possibility is that it may be possible to develop sample preparation, staining, or analysis methods that can partially adjust for the consequences of excessive fluid accumulation, including AIZs artifacts, potentially allowing for inference of neural morphology despite these artifacts. A previous approach to address extracellular space shrinkage in EM images classified the space into sheet-like and tunnel-like regions, then used computational models to redistribute volume between these compartments (Kinney et al., 2013). However, it is highly questionable that approaches designed for controlled shrinkage artifacts could be sufficient for adjustment of postmortem brain EM images, as the AIZ artifacts present are anisotropic, haphazard, and enigmatic. Instead, more complex computational approaches, developed in consideration of the physiology of agonal and postmortem degradation, would likely be required, to the extent that it is feasible at all.

Our light microscopy findings provide complementary data to help interpret the EM results. H&E and immunohistochemical staining revealed relatively well-preserved cellular morphology in samples from the same brains that had substantial changes on EM images. Particularly notable was the integrity of neurofilament staining, which allowed for visualization of axonal and dendritic trajectories across substantial distances despite varying PMIs. This is consistent with previous data showing the robustness of neurofilament antigen staining to extended PMI (Ulfig et al., 1998; Blair et al., 2016; Bouvier et al., 2016). As a result, our findings corroborate previous observations that electron microscopy is more sensitive to postmortem changes than light microscopy (Krassner et al., 2023). One obvious potential explanation for this discrepancy is simply that EM has better resolution and is therefore better able to detect early, minute aspects of tissue decomposition. It is also possible that relative differences in the sensitivity of light microscopy and EM staining procedures to postmortem changes could help to explain the discrepancy. For example, osmium tetroxide, a key staining agent for EM, has been found to bind primarily to unsaturated fatty acids arranged in lipid membranes (Wigglesworth, 1957, 1981; Belazi et al., 2009; Li et al., 2024). Autolysis causes a rapid release and redistribution of membrane lipids, including unsaturated fatty acids, via the activity of phospholipase A2 (van der Vusse et al., 1989; Pueyo et al., 2000). If unsaturated fatty acids in lipid membranes degrade relatively faster than proteins in the PMI, then this would disproportionately affect visualization following standard EM protocols, compared to other methods that target protein antigens. This may help explain why the visualization of unmyelinated axons, which have fewer lipids, is more rapidly lost in postmortem tissue than the visualization of myelinated axons (Liewald et al., 2014). As another possibility, cellular structures might remain partially present during the postmortem period, but their constituent biomolecules could become insufficiently compact for adequate visualization by non-specific morphological stains alone. Immunostaining in particular may be better able to amplify partially intact structures and therefore not be affected as rapidly by the postmortem breakdown of gel-like biomolecular networks as the typical non-specific EM staining methods (Krassner et al., 2023). This possibility would parallel light microscopy studies on postmortem brain tissue showing that immunostaining can sometimes reveal cellular structures not visible with non-specific morphological stains (Monroy-Gómez et al., 2020).

Further research using complementary methods, such as immunoelectron microscopy, correlative light and electron microscopy, or expansion microscopy may help to clarify the reasons for the discrepancy between light and electron microscopy.

Several limitations should be considered when interpreting our findings. First, our sample size of ten brains obviously does not capture the full spectrum of preservation quality variation seen in brain banking. Information about the donors, such as the PMI, may have inaccuracies, which becomes a bigger problem with smaller sample sizes. As a result, our findings will require larger subsequent studies for corroboration. Second, our analysis of the contralateral hemispheres for light and electron microscopy introduces obvious potential confounding factors. Preservation quality, including the quality of perfusion, can clearly vary between the hemispheres, which might affect our results. Third, although formaldehyde alone has been used successfully for fixation in some studies, others recommend the addition of glutaraldehyde (Gonzalez Aguilar and De Robertis, 1963; Westrum and Lund, 1966; Lewis et al., 2019). On initial fixation we used formaldehyde alone without the addition of glutaraldehyde, which may have reduced the ultrastructural quality of our resulting EM images. Fourth, our semi-quantitative grading systems for the EM and immunohistochemically stained images represent a simplified assessment of complex data. Further, our analysis of the light microscopy images was largely qualitative, which renders this more vulnerable to bias. We encourage any interested readers to analyze the raw image data, all of which we have made available, to come to their own conclusions. Lastly, our study focused primarily on 2D ultrastructural features. The implications for 3D connectivity reconstruction remain speculative without more direct volumetric data and analysis. These limitations highlight the need for future studies with larger sample sizes and more direct comparisons via multiple imaging techniques.

## Conclusions

Our study provides insights into the relationship between preservation quality, PMI, and imaging modality in banked brain tissue. We found that the level of ultrastructural preservation required for performing connectomics studies appears to be achievable in some postmortem samples.

These findings suggest that under close to optimal brain banking conditions, postmortem human brain tissue could potentially be suitable for connectomics studies using current EM methods.

However, in at least a subset of EM images from the other samples, our analysis revealed the presence of AIZ artifacts, which may indicate significant challenges for the reliable tracing of neural processes. These AIZs artifacts may result from true structural degradation prior to fixation, inadequate visualization of thin axons and other cellular components, or both. Notably, our light microscopy data from these same brains indicates that many cellular structures remain intact and potentially traceable at this level of resolution via neurofilament staining. This suggests a discrepancy between preservation quality as observed via different imaging modalities. To advance connectomics research using banked brain tissue, further methodological development may be valuable. Current approaches can only reliably visualize key ultrastructural features in a small subset of banked brain samples, constraining the possible sample sizes. It is not yet clear if these are the true biological limits to ultrastructural visualization. Future research could focus on optimizing fixation protocols, staining techniques, and computational methods specifically for postmortem brain tissue. Such research may be critical for realizing the full potential of human brain connectomics as a tool for understanding neural circuit organization in both health and disease.

## Abbreviations

EM: Electron Microscopy
GFAP: Glial Fibrillary Acidic Protein
h: Hour
H&E: Hematoxylin and Eosin
IRR: Inter-Rater Reliability
LM: Light Microscopy
NBF: Neutral Buffered Formalin
PMI: Postmortem Interval
WSI: Whole Slide Image.

## Author contributions

A.K., M.G., K.F., J.C., and A.T.M. conceptualized the article. A.S. and W.J. performed electron microscopy experiments. E.T. and C.D.S. performed light microscopy experiments. A.K., M.G., and A.T.M. performed data analysis. A.T.M. wrote the initial draft of the manuscript. All authors reviewed the manuscript. All authors approved the final manuscript.

## Acknowledgements

We would like to acknowledge the technical assistance of Laura Paredes and Alexander Parra, and we would like to thank Kenneth Hayworth for helpful personal communications on this topic. We acknowledge the Neuropathology Brain Bank & Research CoRE at the Icahn School of Medicine at Mount Sinai for their histology and tissue processing services. Electron microscopy tissue preparation and imaging were performed at The Microscopy and Advanced Bioimaging CoRE at the Icahn School of Medicine at Mount Sinai. The Icahn School of Medicine at Mount Sinai provided access to library resources.

## Funding

This work was supported by the Rainwater Charitable Foundation as well as NIH grants P30 AG066514, K01 AG070326, RF1 AG062348, P30 AG066514, RF1 NS095252, U54 NS115266, and RF1 MH128969. The funders had no role in the design of the study or in the collection or interpretation of the data.

## Conflict of interest

Alicia Keberle, Macy Garrood, and Andrew McKenzie are employees of Oregon Brain Preservation, a non-profit brain preservation organization. Andrew McKenzie is a director of Apex Neuroscience, a non-profit research organization.

## Data availability

Code used for data analysis is available here: https://github.com/andymckenzie/Ultrastructure_quality_manuscript. All raw image data can be accessed in a public repository on Zenodo, available here: https://zenodo.org/communities/evaluatingultrastructuralpreservationqualityinbankedbraintissue/

## Notes

https://zenodo.org/communities/evaluatingultrastructuralpreservationqualityinbankedbraintissue/

https://github.com/andymckenzie/Ultrastructure_quality_manuscript

